# Nogo-A regulates the fate of human dental pulp stem cells towards osteogenic, adipogenic, and neurogenic differentiation

**DOI:** 10.1101/2022.09.01.506142

**Authors:** Chai Foong Lai, Juliet Shen, Anamaria Balic, Pierfrancesco Pagella, Martin E. Schwab, Thimios A. Mitsiadis

**Author notes:** **Corresponding author:** Thimios A. Mitsiadis, Institute of Oral Biology, Center of Dental Medicine, University of Zurich., Plattenstrasse 11, CH8032, Zurich, Switzerland.

## Abstract

Human teeth are highly innervated organs that contain a variety of mesenchymal stem cell populations that could be used for cell-based regenerative therapies. Specific molecules are often used in these treatments to favorably modulate stem cells function and fate. Nogo-A, a key regulator of neuronal growth and differentiation, is already used in clinical tissue regeneration trials. While the functions of Nogo-A in neuronal tissues are extensively explored, its role in teeth still remains unknown. In this work, we first immunohistochemically analyzed the distribution of Nogo-A protein in the dental pulp of human teeth. Nogo-A is localized in a variety of cellular and structural components of the dental pulp, including odontoblasts, fibroblasts, neurons and vessels. We also cross-examined *Nogo* expression in the various pulp cell clusters in a single cell RNA sequencing dataset of human dental pulp, which showed high levels of expression in all cell clusters, including that of stem cells. We then assessed the role of Nogo-A on the fate of human dental pulp stem cells and their differentiation capacity *in vitro*. Using immunostaining, Alizarin Red S and Oil Red O staining we showed that Nogo-A delayed the differentiation of cultured dental pulp stem cells towards the osteogenic, adipogenic and neurogenic lineages, while addition of the blocking anti-Nogo-A antibody had opposite effects. These results were further confirmed by qRT-PCR, which demonstrated overexpression of genes involved in osteogenic (*RUNX2, ALP, SP7/OSX*), adipogenic (*PPAR-γ2, LPL*) and neurogenic (*DCX, TUBB3, NEFL*) differentiation in presence of the anti-Nogo-A antibody. Conversely, the osteogenic and adipogenic genes were downregulated by Nogo-A. Taken together, our results show that the functions of Nogo-A are not restricted to neuronal cells, but are extended to other cell populations, including dental pulp stem cells. We show that Nogo-A regulates their fates towards osteogenic, adipogenic and neurogenic differentiation, thus indicating its potential use in the clinics.

## Introduction

Tissue homeostasis and regeneration are ensured by stem cells, which have the capability to self-renew, proliferate and differentiate towards distinctive cell lineages ^1–3^. In the human body, diverse stem cell populations reside in specific anatomic locations, called stem cell niches, which support stem cell maintenance and influence their fates ^4–10^. Stem cells administered to injured tissues and organs support their regeneration since they can differentiate towards distinctive tissue-specific cell types, a process conditioned by the local microenvironment ^1,4,8,9,11–15^. Human teeth are characterized by several distinct mesenchymal stem cell populations, including those found in the pulp of human adult teeth and/or exfoliated deciduous teeth, the apical papilla, the dental follicle and the periodontal ligament ^12,16–18^. These stem cell populations vary in their expression of stem cell-related genes, their ability to differentiate into distinctive cell lineages, as well as the composition of the niche in which they reside ^4,8,10,11,13,19,20^. Dental mesenchymal stem cells can be easily isolated from the pulp of healthy human adult teeth, which is a heterogeneous tissue composed of odontoblasts, fibroblasts, immune cells, nerve ensheathing cells, vascular cells and various mesenchymal stem cell populations with different stages of lineage commitment ^11–13,18,21^. The hallmark of human dental pulp stem cells (hDPSCs) is the maintenance of tooth homeostasis and the regeneration of the dentin-pulp complex upon injury ^18,22^–^261^–^6^. We and others have previously demonstrated the remarkable plasticity of hDPSCs and their ability to differentiate into various cell lineages, including osteogenic, adipogenic, vascular, myogenic and neurogenic lineages ^13,27–37^, thus indicating that these cells represent a precious source of stem cells with great clinical potential for tissue regenerative approaches ^12,23,26,30,38–42^. Indeed, few clinical studies have already shown the ability of hDPSCs to regenerate pulp tissue after pulpectomy, as well as their potential use toward bone regeneration *in vivo* ^22,26,43–45^.

Current regenerative treatments are also based on exploiting molecules with specific cellular functions that modulate the differentiation potential of the stem cells ^39,46,47^. One of these molecules used in tissue repair is the 200 kDa Nogo-A protein, which acts as an inhibitor of neurite outgrowth and plasticity ^48–51^. Nogo-A exerts its signaling functions upon interaction with receptor complexes, including the S1PR2 receptor ^50^, and the NgR1, LINGO-1 (immunoglobulin-like domain-containing protein-1), p75NTR (p75 neurotrophin receptor) and TROY (TNF receptor superfamily member 19) containing receptor complex ^52–58^. Apart from its well-known neuronal functions, Nogo-A may act as a proliferative, differentiation, survival and apoptotic agent in non-neuronal cell populations ^59–62^. For example, various studies have involved Nogo-A in the regulation of angiogenesis and vascular remodeling, macrophage activation during inflammatory processes, and stem cell behavior ^63-717-15^. These findings, together with successfully completed ^72^ or ongoing clinical trials in spinal cord injured patients illustrate the important role of Nogo-A in a variety of biological processes, including tissue regeneration.

Surprisingly, there is yet no information on the function of Nogo-A in the highly innervated dental tissues, despite its reported expression in human teeth ^73^. In this study we examined the distribution of the Nogo-A protein in the dental pulp of human teeth and we assessed its role and effects on the plasticity of hDPSCs by analyzing their differentiation potential *in vitro*. These studies were complemented by gene expression analyses of specific stem cell, osteogenic, adipogenic and neuronal markers. Our results show that Nogo-A plays an important role in stem cell fate choices and differentiation events, and point to Nogo-A as a suitable candidate molecule for future regenerative therapies in the clinics.

## Materials and Methods

### Data and code availability

Data can be found at GEO; Access number: GSE161267. All code is publicly available at: https://github.com/TheMoorLab/Tooth. Detailed single cell RNA sequencing data is previously described ^11^.

### Collection of human tissues and cells

Healthy human teeth were obtained from anonymous patients at the Center of Dental Medicine, of University of Zurich. The Ethics Committee of the Canton of Zurich has approved the project (reference number 2012-0588) and the patients gave their written informed consent. The teeth were extracted by dentists at the Clinic of Cranio-Maxillofacial and Oral Surgery Department at the Center of Dental Medicine of the University of Zurich. All procedures were implemented following the current guidelines. Human dental pulp cells were isolated from the dental pulp of extracted wisdom teeth of healthy patients as previously described ^11^. Briefly, the dental pulps were gently removed with forceps, minced with scalpels, rinsed with PBS and finally were enzymatically digested for 1 hr at 37° C in a solution of Collagenase (3 mg/mL; Life Technologies BV, Zug ZG, Switzerland) and Dispase (4 mg/mL; Sigma-Aldrich Chemie GmbH, Buchs SG, Switzerland).

### Cell cultures and differentiation assays

Dental pulp cells were seeded at a density of 30,000 cell/cm^2^ using Dulbecco’s Modified Eagle Medium / Nutrient Mixture F-12 (DMEM/F12) medium containing GlutaMAX (Gibco, 31331-028), supplemented with 10% Fetal Bovine Serum (FBS; Sigma, F0804) and 1% penicillin/streptomycin (Gibco, 15070-063). The cultured cells were incubated at 37° C in a humidified atmosphere of 5% CO2, 95% air, and the culture medium was changed every other day. The cells were treated with either anti-Nogo-A monoclonal antibody (10 μg/ml) or Nogo-A Δ20 protein (20 nM) for the duration of the culture. Osteogenesis, adipogenesis and neurogenesis-induced media were applied when the cells reached 80-90 % confluence ^29,35^.

For the induction of osteogenenesis the cells were cultured in DMEM medium (Gibco, 21885-025), supplemented with 10% FBS (Sigma, F0804) and 1% penicillin/streptomycin (Gibco, 15070-063), ascorbic acid (200 μM; Sigma, A4544), ß-glycerophosphate (10 mM; Sigma, G9422) and dexamethasone (10 nM; Sigma, D4902). For the adipogenenic assay, confluent cells were cultured in DMEM medium, 10% FBS, 1% penicillin/streptomycin, dexamethasone (1 μM), 3-isobutyl-1-methylxanthine (0.5 mM; Sigma, 28822-58-4), indomethacin (200 μM; Sigma, I7378) and insulin (10 μM; Sigma, I9278). For the induction of neurogenesis cells were cultured in presence of Neurobasal A medium (Gibco, 21103-049), 1% penicillin/streptomycin), 1% B27 (Gibco, 17504-044), Epidermal Growth Factor (EGF; 20 ng/ml; PeproTech, AF-100-15) and Fibroblast Growth Factor (FGF; 40 ng/ml; PeproTech, 100-17A). The morphology of the cells was analyzed after 7, 14, and 21 days of induction. The experimental plan is provided in Table 1.

**Table 1.**
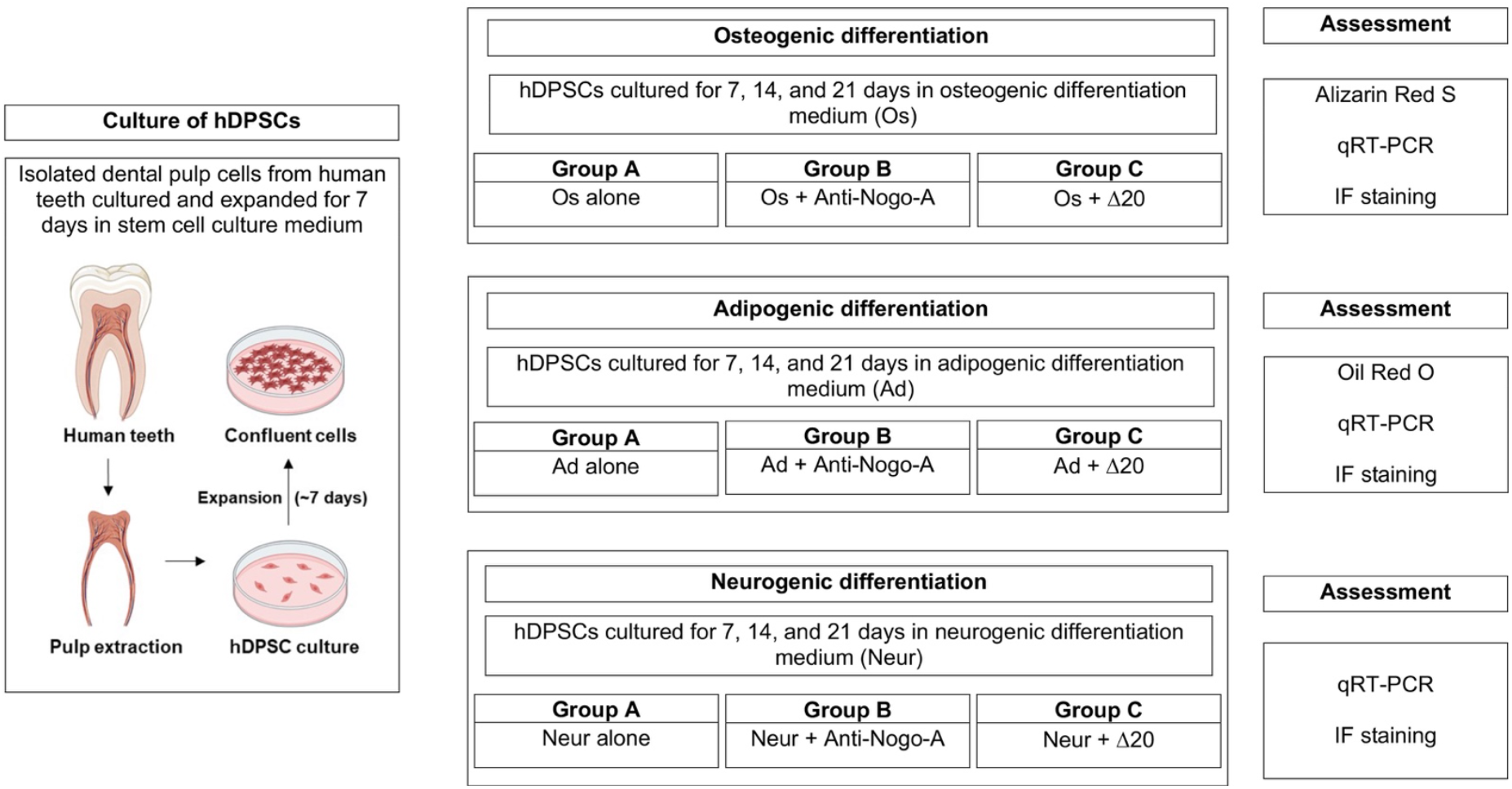
Experimental plan for analyzing the effects of Nogo-A (Nogo-A Δ20 fragment) and anti-Nogo-A antibody on the osteogenic, adipogenic, and neurogenic differentiation of human dental pulp stem cells.

### Staining

#### Immunohistochemistry

Immunoperoxidase staining on paraffin sections was performed as previously described ^74,75^. Briefly, 5 μm sections were deparaffinized, exposed to a 0.3 % solution of hydrogen peroxide in methanol, and then incubated overnight at 4°C in a humid atmosphere either with the mouse monoclonal anti-Nogo-A antibody ^76^ diluted 1:100 in PBS / 0.2% BSA. Peroxidase was revealed by incubation with 3-amino-9-ethylcarbazole (AEC) reaction solutions. After staining the sections were mounted with Eukit. In control sections the primary antibodies were omitted. Immunohistochemistry on cultured dental pulp cells was performed after 4% paraformaldehyde (PFA) in PBS fixation, followed by permeabilization of cells for 15 min with 0.5 % Triton X-100 in PBS. Fixed and permeabilized samples were then incubated with the primary antibodies overnight at 4°C, and the signal was detected by AEC. Upon staining the slides were mounted with Aquamount (BDH, Gurr, England). Controls were performed by incubations without primary antibodies.

#### Immunofluorescence Staining and Imaging

Polyclonal anti-Collagen type-I antibody (ab34710, Abcam), rabbit anti-FABP4 monoclonal antibody (EPR3579, Abcam), rabbit anti-Neurofilament light chain (NFL; NB300-131, Novus biology), rabbit anti-Tubulin Beta-III (TUBB3; ab18207, Abcam), and alexa-488-conjugated anti-rabbit IgG (A21206, Abcam) antibodies were used. The cells on IBIDI μ-Slide 8-well chamber slide were fixed with 4% PFA for 10 min, after which the fixative was removed and cells were washed twice with PBS. Next, the cells were permeabilized with PBST (0.1% Triton X-100 in PBS) for 10 min at room temperature (RT). Slides were then washed thrice with PBST (5 min each). Non-specific binding was blocked by incubation with 5% BSA in PBST for 1 hr at RT. The cells were incubated in primary antibodies for 1 hr at RT. Slides were washed three times with PBST before incubation with Alexa Fluor® 488 donkey anti-rabbit IgG secondary antibody (Invitrogen, Carlsbad, CA, USA), for 1 hr at RT. Nuclei were stained with DAPI (1ug/mL) for 10 min after which the chamber slides were mounted in ProLong™ Diamond Antifade Mountant (Invitrogen, P36961). All images were acquired using Leica Stellaris confocal microscope.

#### Alizarin Red S Staining

The extend of mineralization was analyzed in cultures incubated under osteogenic conditions, at different time points (7, 14, and 21 days). Cultures were washed twice with PBS and fixed with 4% PFA for 10 min at RT. Fixed cultures were rinsed with distilled water and incubated with Alizarin Red S (Sigma-Aldrich) for 15 min at RT. Excess Alizarin Red was removed with three washes of distilled water.

#### Oil Red O Staining

The work solution was made prior to staining by mixing the Oil Red O stock solution (0.5% in isopropanol) with distilled water in 3:2 ratio, and leaving it for 1 hr at RT followed by filtration through 0.22 μm pore size filters. hDPSCs cells were fixed in 12-well plates with 4% PFA in PBS for 20 min. Fixed cells were stained with Oil Red O for 20 min and after rinsing several times with PBS they were observed through an inverted microscope. The intracellular Oil Red O staining was eluted upon incubation with isopropanol for 15 min. The optical density was assessed using the ELISA reader at 500 nm.

### RNA Extraction, Reverse Transcription and Quantitative Real-Time PCR (qRT-PCR)

Total RNA was extracted from the cells using TRIzol reagent (Ambion, 15596026) according to the manufacturer’s instructions. The reverse transcription of the RNA was performed by using the iScript™ cDNA synthesis Kit (Cat. No. 1706691; Bio-Rad Laboratories AG, Hercules CA, USA). Relative mRNA expression levels were evaluated using the SYBR Green method. The quantitative 3-step real-time PCR was performed by the Eco Real-Time PCR system (Illumina Inc., San Diego CA, USA) and CFX Connect (Bio-Rad Laboratories AG, Hercules CA, USA). Expression levels of *GAPDH, CD90/THY1, CD105, RUNX2, ALP, OSX, LPL, PPARγ2, βIII-Tubulin, Neurofilament-L*, and *DCX* were analyzed. The sequences of primers are supplied in Table 2. All primers were synthesized by Microsynth (Microsynth AG, Balgach, Switzerland). All of the reactions were run in triplicates. After the reactions were realized, the Cq values were established by setting a fixed threshold. All measurements were normalized to GAPDH and the relative changes in target gene expression between the control and treatment groups were measured using the 2-^ΔΔ^CT method ^77^.

**Table 2.**
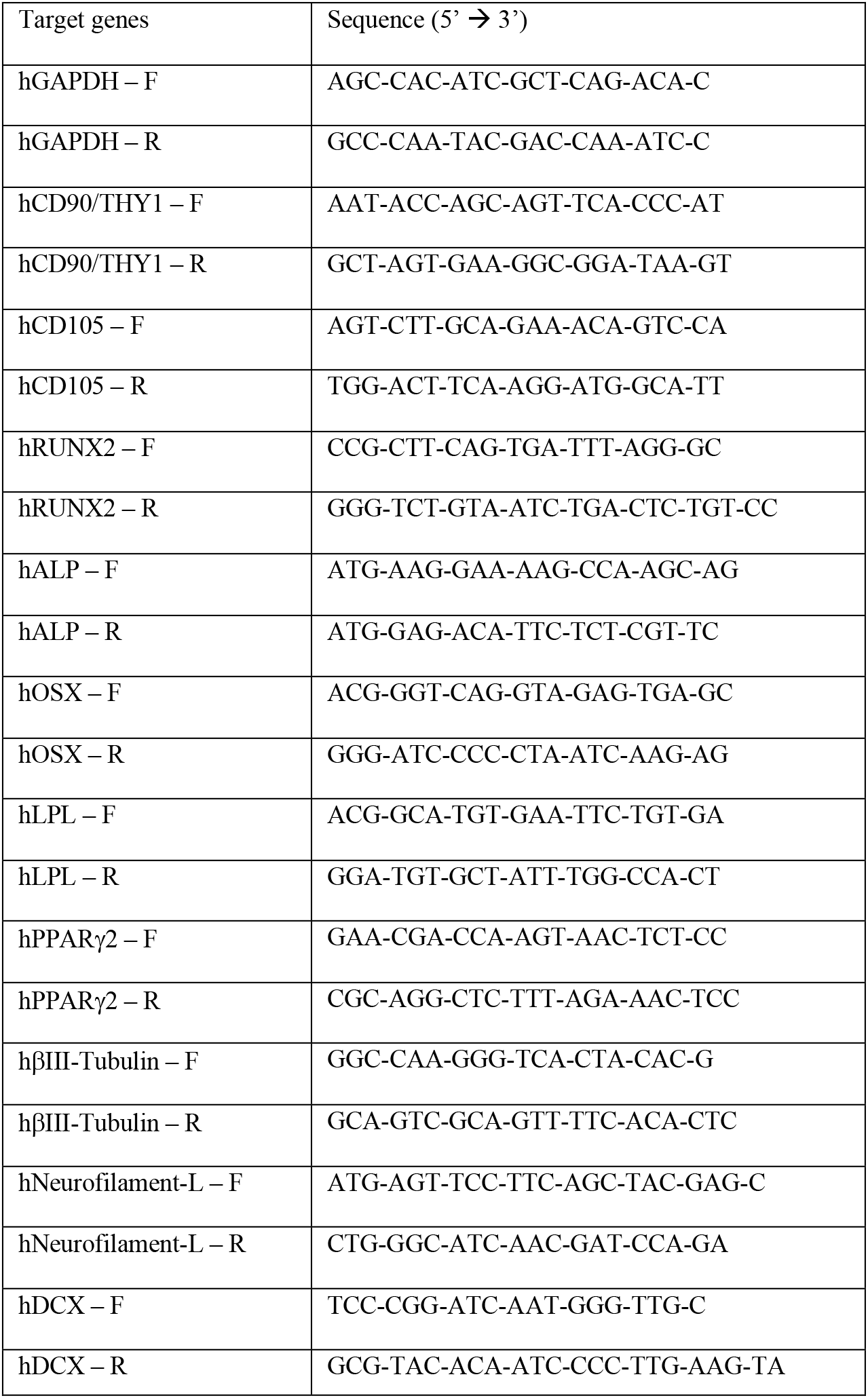
Gene sequences used for qRT-PCR for the analysis of the osteogenic, adipogenic and neurogenic potential of human dental pulp stem cells.

### Statistical analysis

All qRT-PCR data were analyzed with Student’s t-test. P values < 0.05 were considered statistically significant. Statistical analysis was accomplished with Microsoft Excel Software (Microsoft Corporation, USA, Version 16.55).

## Results

### Localization of the Nogo-A protein in human dental pulp cells

We analyzed the distribution of Nogo-A in the dental pulp of healthy permanent human teeth (Fig. 1A-C), and found that this molecule was abundantly expressed within this tissue (Fig. 1D-H). Nogo-A immunoreactivity was observed in various dental pulp cell populations, including odontoblasts, fibroblasts, nerves and vessels, albeit at different intensities (Fig. 1D-H). Intense staining was detected in the nerves and blood vessels (Fig. 1B-H), while strong staining was also observed in odontoblasts (Fig. 1B, D, E). Immunohistochemical (Fig. 1I) and immunofluorescent (Fig. 1J) staining was also detected in dissociated human dental pulp cells that were cultured for 5 days *in vitro*. Immunostaining was not observed in control sections, in which the Nogo-A antibody was omitted (data not shown).

**Figure 1.**
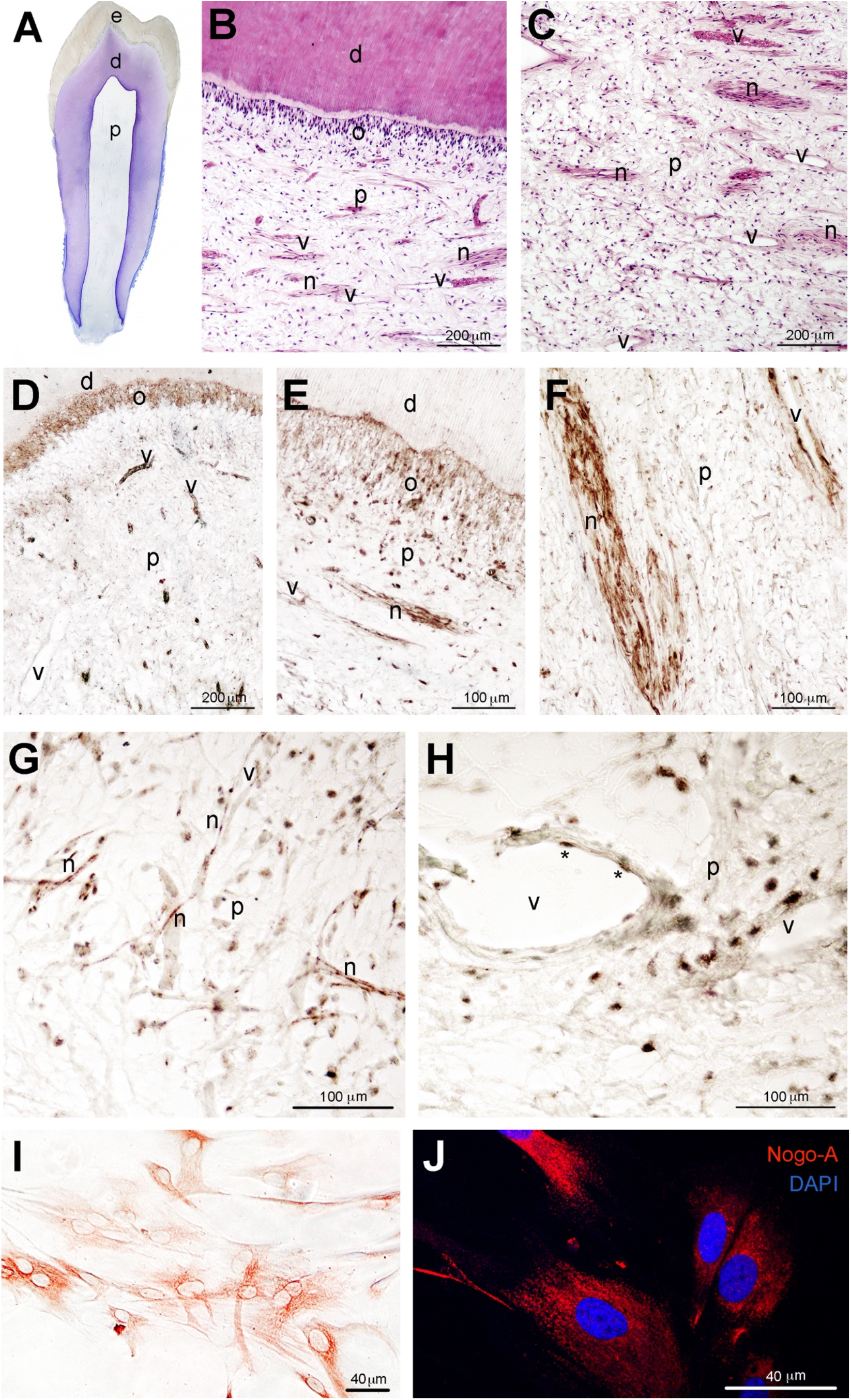
Nogo-A protein distribution in the dental pulp of human adult teeth and in cultured human dental pulp stem cells (hDPSCs). (A) Hematoxylin and Eosin (H&E) stained histological section of a human first premolar. (B, C) Histological sections of human adult teeth stained with H&E, showing the various components of the dentin-pulp complex. (D-H) Immunostaining showing the distribution of Nogo-A (brown color) in odontoblasts (D, E), pulp fibroblasts (E, G, H), vessels (D, F, H) and nerves (E, F, G) of human teeth. Asterisks in (H) indicate endothelial cells. (I) Nogo-A immunostaining (brown/red color) in cultured hDPSCs. (J) Nogo-A immunofluorescence staining (red color) in hDPSCs. DAPI (blue color) marks the nuclei of the cells. Abbreviations: d, dentin; e, enamel; n, nerves; o, odontoblasts; p, dental pulp; v, vessels.

We have performed recently a detailed single cell RNA sequencing analysis of the human dental pulp and periodontium ^11^. We have interrogated this dataset to determine the number of cells expressing *NOGO* (*RTN4*) and the *NOGO*-related receptors (*LINGO1, RTN4R, S1PR2, TNFRSF19, NGFR*) in the stem cell compartment of the pulp (DPSC compartment; blue color in Fig. 2A), which represents the 12 % of the dental pulp tissue (4’674 cells in a total number of 32’378 cells; Fig. 2A, H) and highly express the *FRZB, NOTCH3, THY1*, and *MYH* genes ^11^. In the DPSC compartment, 1’523 cells expressed *RTN4* in a total number of 4’674 cells, which represent approximately the 33 % of all DPSCs (Fig. 2B, H). Cells expressing *LINGO1, RTN4R, S1PR2, TNFRSF19, NGFR* were less abundant in the DPSC compartment (Fig. 2C-H).

**Figure 2.**
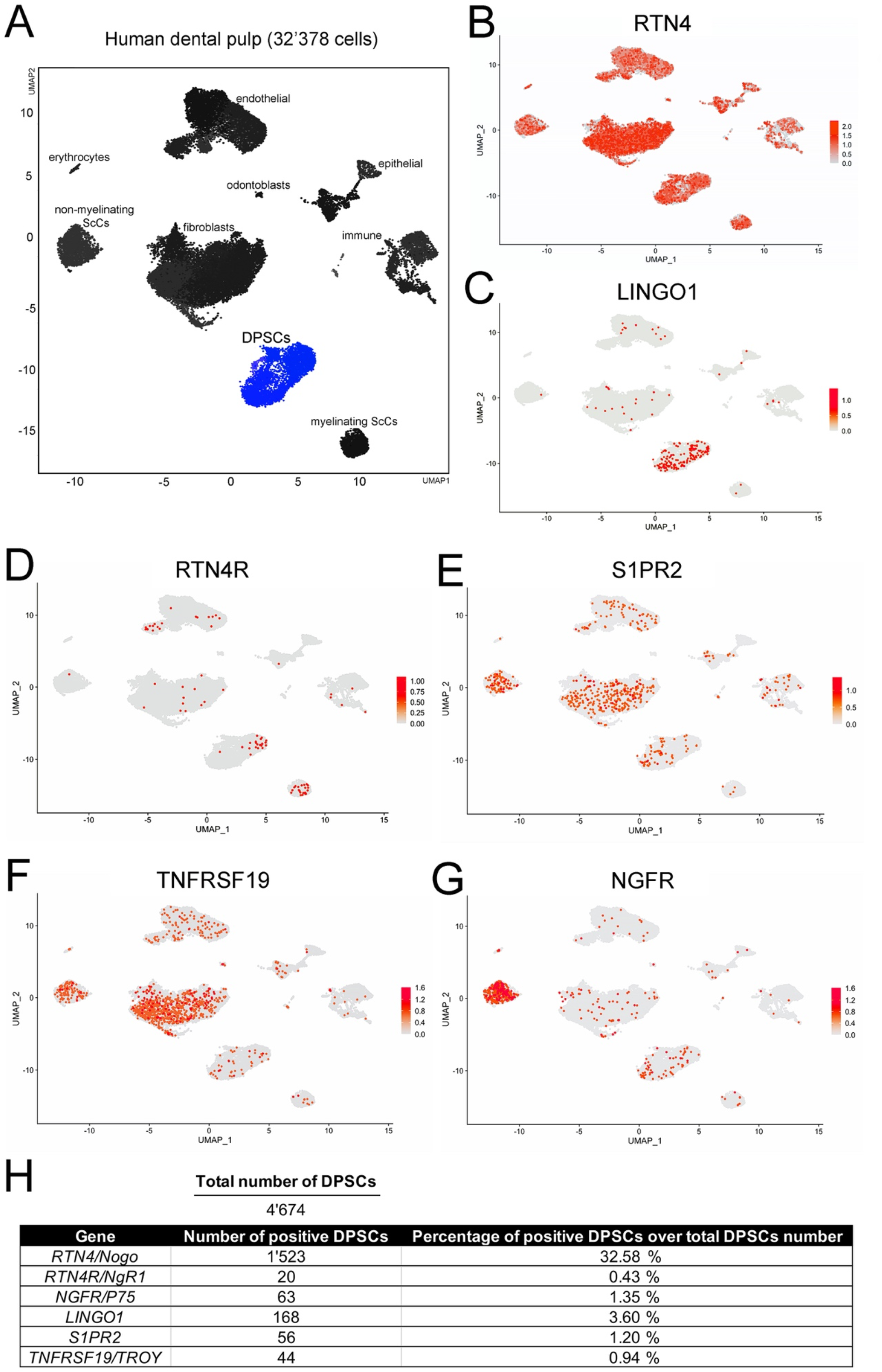
Feature plots showing the expression of *NOGO* and *NOGO-related* receptors genes in the dental pulp stem cell compartment of human adult teeth. (A) Specific cell clusters within the dental pulp. In blue the DPSCs cluster. (B-G) Cells expressing *RTN4 (NOGO;* B), LINGO1 (C), RTN4R (D), S1PR2 (E), TNFRF19 (F), and NGFR (G) in all dental pulp compartments. Expression in individual cells is represented by red dots. (H) Quantitative details of the *NOGO* and *NOGO*-related receptors genes within DPSCs cluster.

### Nogo-A effects on the stemness of hDPSCs

We first analyzed by RT-qPCR the relative expression of stem cell markers (THY1/CD90 and END/CD105) in hDPSCs cultured in DMEM maintenance medium alone (group A), supplemented with the anti-Nogo-A antibody (group B), and supplemented with the Nogo-A Δ20 protein (group C) for 7 days (Fig. 3). The expression of *THY1* was increased in hDPSCs from group C, while expression was decreased in hDPSCs from group B (Fig. 3A). In contrast, the expression of *CD105* was increased in hDPSCs from the group B, while expression was unaffected in hDPSCs of the group C (Fig. 3B). These results suggest that Nogo-A is critical for maintaining the stemness of hDPSCs.

**Figure 3.**
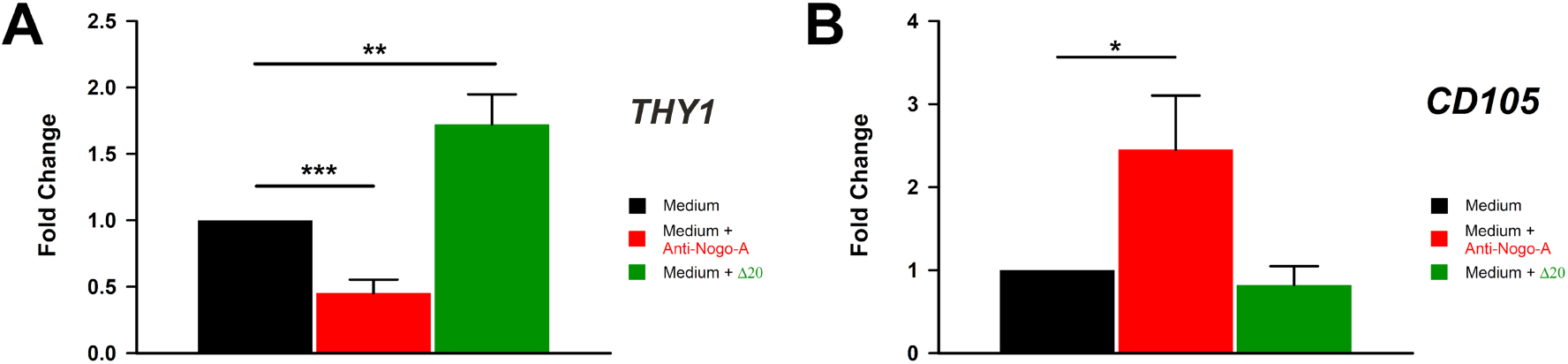
Effects the Nogo-A protein and anti-Nogo-A antibody on the expression of *THY1/CD90* and *END/CD105* upon culture of hDPSCs in DMEM maintenance medium. qRT-PCR analysis showing *THY1/CD90* (A) and *END/CD105* (B) relative expression levels in hDPSCs cultured in DMEM maintenance medium alone (black columns), in medium supplemented with the anti-Nogo-A-antibody (red colored columns), and medium supplemented with the Nogo-A Δ20 protein (green colored columns) for 7 days. Values, normalized to Glyceraldaeyde-3-Phosphate Dehydrogenase (GAPDH), shown are the mean ± SD of triplicates. Statistical analysis used Student’s t-test (significant difference at *, p<0.02; **, p<0.002; ***, p<0.005).

### Nogo-A effects on the *in vitro* osteogenic, adipogenic, and neurogenic differentiation potential of hDPSCs

Thereafter, the effects of Nogo-A on the multilineage differentiation potential of hDPSCs were assessed in cultures induced for osteogenic, adipogenic and neurogenic differentiation.

#### The anti-Nogo-A antibody accelerates the osteogenic differentiation of hDPSCs

hDPSCs obtained from healthy permanent teeth were cultured in osteogenic medium alone (control, group A), in osteogenic medium supplemented with the anti-Nogo-A antibody (group B), and in osteogenic medium supplemented with the active Nogo-A protein fragment Δ20 (group C). The osteogenic potential of hDPSCs in the three groups was assessed 7, 14 and 21 days upon culture by their ability to form extracellular calcium deposits (mineralized nodules), which were detected by Alizarin Red S staining (Fig. 4). hDPSCs from group A started to form mineralized nodules between 14 and 21 days of culture (Fig. 4E, H). hDPSCs from group B displayed early signs of osteogenesis and formed prematurely mineralized nodules as compared to hDPSCs of groups A and C (compare Figs 4B, E, H and 4D, G, J with Fig. 4C, F, I). In group A, the mineralized nodules were first observed at 14 days of culture (Fig. 4F). Their size and the extent of mineralization significantly and continuously increased up to the end of the culture period (Fig. 4I). In contrast, hDPSCs from group C delayed the onset of mineralization and reduced the overall mineralization capacity as compared to hDPSCs of groups A and B (compare Figs 4B, E, H and 4C, F, I with Fig. 4D, G, J). These observations were further confirmed by quantification of Alizarin Red S incorporated in the mineralized nodules, which showed its significant increase at 14 and 21 days in group B as compared to groups A and C (Fig. 4A). Conversely, hDPSCs from group C demonstrated significantly lower level of incorporated Alizarin Red S as compared to hDPSCs of groups A and B (Fig. 4A). Increased collagen type-I immunostaining was also observed in hDPSCs from group B (Fig. 4L) as compared to hDPSCs of groups A (Fig. 4K) and C (Fig. 4M).

**Figure 4.**
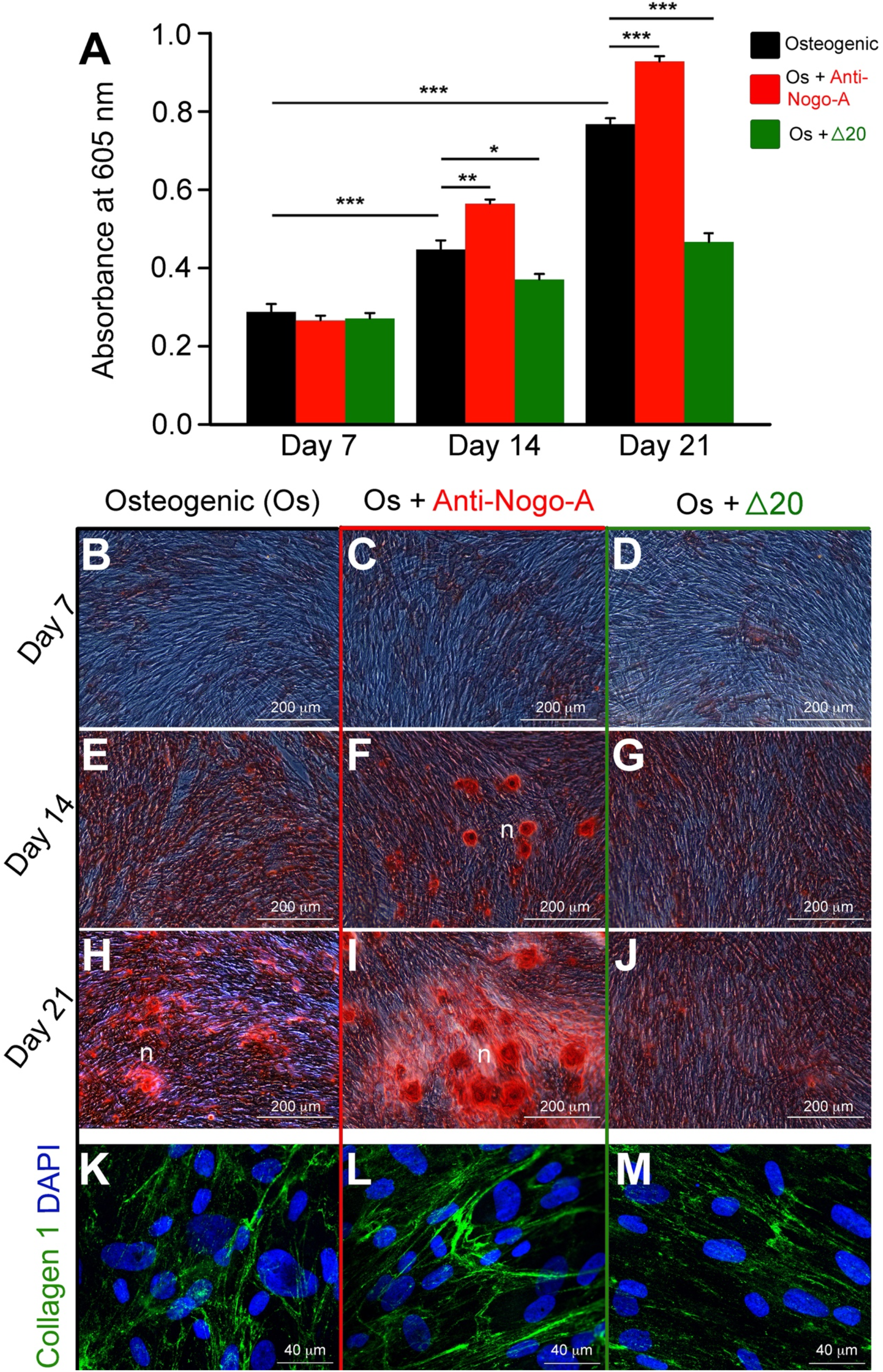
Effects of the Nogo-A protein (Nogo-A Δ20 fragment) and anti-Nogo-A antibody on the osteogenic differentiation of hDPSCs. (A) Quantification of Alizarin Red S staining in hDPSCs cultured in osteogenic medium (Os) alone (black colored columns), in osteogenic medium supplemented with the anti-Nogo-A antibody (red colored columns), and in osteogenic medium supplemented with the Nogo-A Δ20 protein (green colored columns) for 7, 14 and 21 days. Four independent biological replicates were used. The bars represent the means ± standard errors. Asterisks above the error bars denote significant differences (*, p < 0.05; **, p<0.01; ***, p<0.001) by one-way analysis of variance (ANOVA). (B-J) Alizarin Red S staining indicating the formation of mineralized nodules by hDPSCs cultured in the osteogenic medium alone (B, E, H), in osteogenic medium supplemented with the anti-Nogo-A antibody (C, F, I), and in osteogenic medium supplemented with the Nogo-A Δ20 protein (D, G, J) for 7, 14 and 21 days. (K-M) Collagen type-I immunofluorescence staining (green color) in hDPSCs cultured for 21 days in the osteogenic medium alone (K), in osteogenic medium supplemented with the anti-Nogo-A antibody (L), and in osteogenic medium supplemented with the Nogo-A Δ20 protein (M). DAPI (blue color) marks the cell nuclei. Abbreviation: n, mineralization nodules. Scale bars are as indicated.

We next analyzed by RT-qPCR the expression of stem cell (THY1/CD90 and END/CD105) and osteogenenesis-related (RUNX2, ALP and OSX) markers in these cultures (Fig. 5). After 7 days of culture, the expression of *THY1/CD90* was significantly decreased in hDPSCs from group B, and it continuously declined until the end of culture (Fig. 5A). In contrast, hDPSCs from group B significantly increased the expression of *END/CD105* (Fig. 5B). *THY1/CD90* downregulation in hDPSCs from group B was accompanied by a significant increase of *RUNX2* expression, indicating the initiation of osteogenic differentiation already at day 7 (Fig. 5C). A similar upregulation was observed in the expression of *ALP* (Fig. 5D) and *SP7/OSX* (Fig. 5E) after 14 and 21 days of culture in hDPSCs from group B, which marked their ongoing osteogenic differentiation. In contrast, the expression of *THY1* was significantly increased and maintained high throughout the culture period in hDPSCs from group C (Fig. 5A). *THY1* upregulation was associated with a considerable downregulation of *RUNX2, ALP* and *OSX* expression in hDPSCs from group C as compared to hDPSCs from groups A and B (Fig. 5C-E). These results suggest that Nogo-A is critical for maintaining the stemness of hDPSCs and inhibiting their differentiation into osteoblasts.

**Figure 5.**
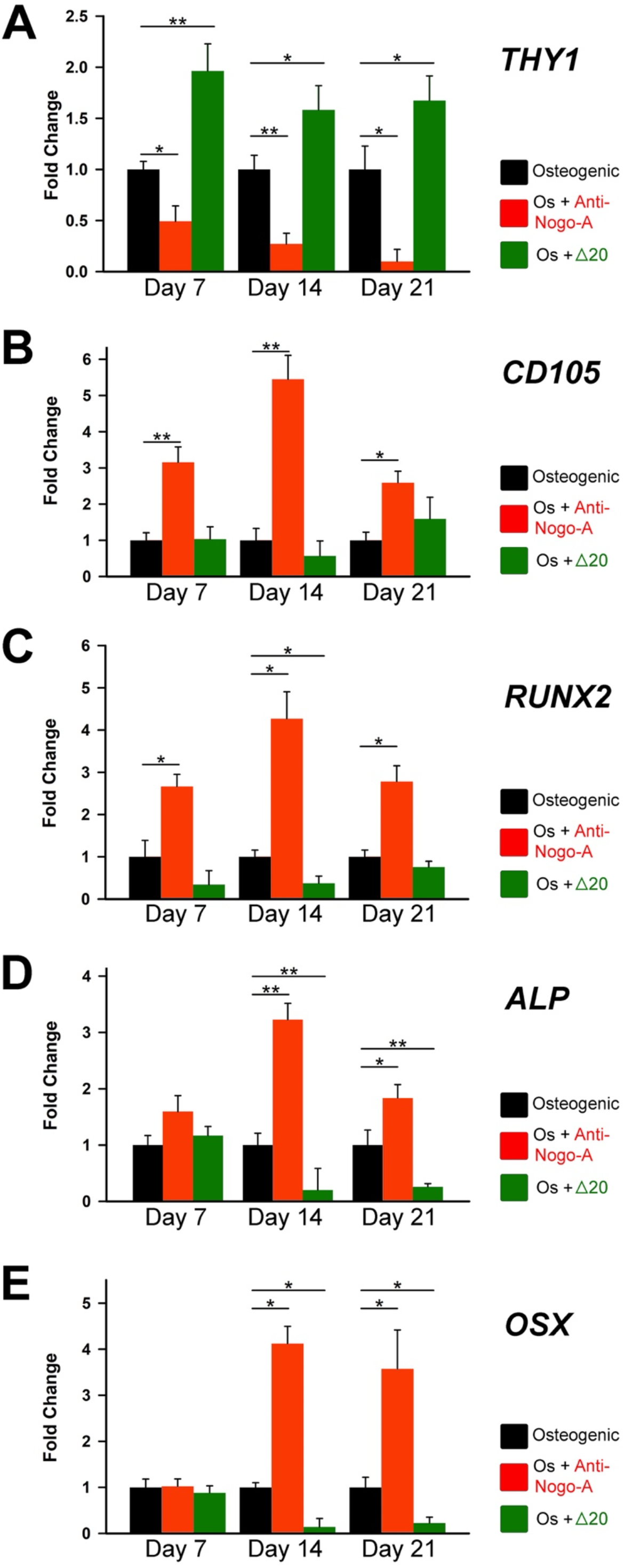
Effects the Nogo-A protein and anti-Nogo-A antibody on the expression of genes during hDPSCs osteogenic differentiation. qRT-PCR analysis showing *THY1/CD90* (A), *END/CD105* (B), *RUNX2* (C), *ALP* (D), and *SP7/OSX* (E) expression levels in hDPSCs cultured in osteogenic medium (Os) alone (black columns), in osteogenic medium supplemented with the anti-Nogo-A-antibody (red colored columns), and osteogenic medium supplemented with the Nogo-A Δ20 protein (green colored columns) for 7, 14 and 21 days. Values, normalized to Glyceraldaeyde-3-Phosphate Dehydrogenase (GAPDH), shown are the mean ± SD of triplicates. Statistical analysis used Student’s t-test (significant difference at *, p<0.05; **, p<0.01).

#### The anti-Nogo-A antibody accelerates the adipogenic differentiation of hDPCSs

hDPSCs can differentiate towards adipocytes if cultured at high cell density in presence of 10% FBS, Dexamethasone, Isobutyl-1-Methylxanthine, Indomethacin and Insulin. We tested the *in vitro* adipogenic differentiation potential of hDPSCs in adipogenic medium alone (control, group A), in adipogenic medium supplemented with the anti-Nogo-A antibody (group B), and in adipogenic medium supplemented with the Nogo-A Δ20 protein (group C), after 7, 14 and 21 days of culture (Fig. 6). The adipogenic differentiation of hDPSCs in the three groups was assessed by their capacity to form and accumulate intracellularly lipid droplets, which were identified by Oil Red O staining (Fig. 6) ^29^. Lipid globules in hDPSCs from group A were first detected around day 14 and their number was increased by the end of the culture period (day 21) (Fig. 6B, E, H). Oil Red O staining in group B hDPSCs was significantly higher at days 14 and 21 as compared to hDPSCs of group A (compare Fig. 6C, F, I with Fig. 6B, E, H). The level of Oil Red O staining in hDPSCs from group C was comparable to group A (compare Fig. 6D, G, J with Fig. 6B, E, H). Oil Red O staining quantification in hDPSCs showed a significant increase at all time points in group B as compared to groups A and C (Fig. 6A).

**Figure 6.**
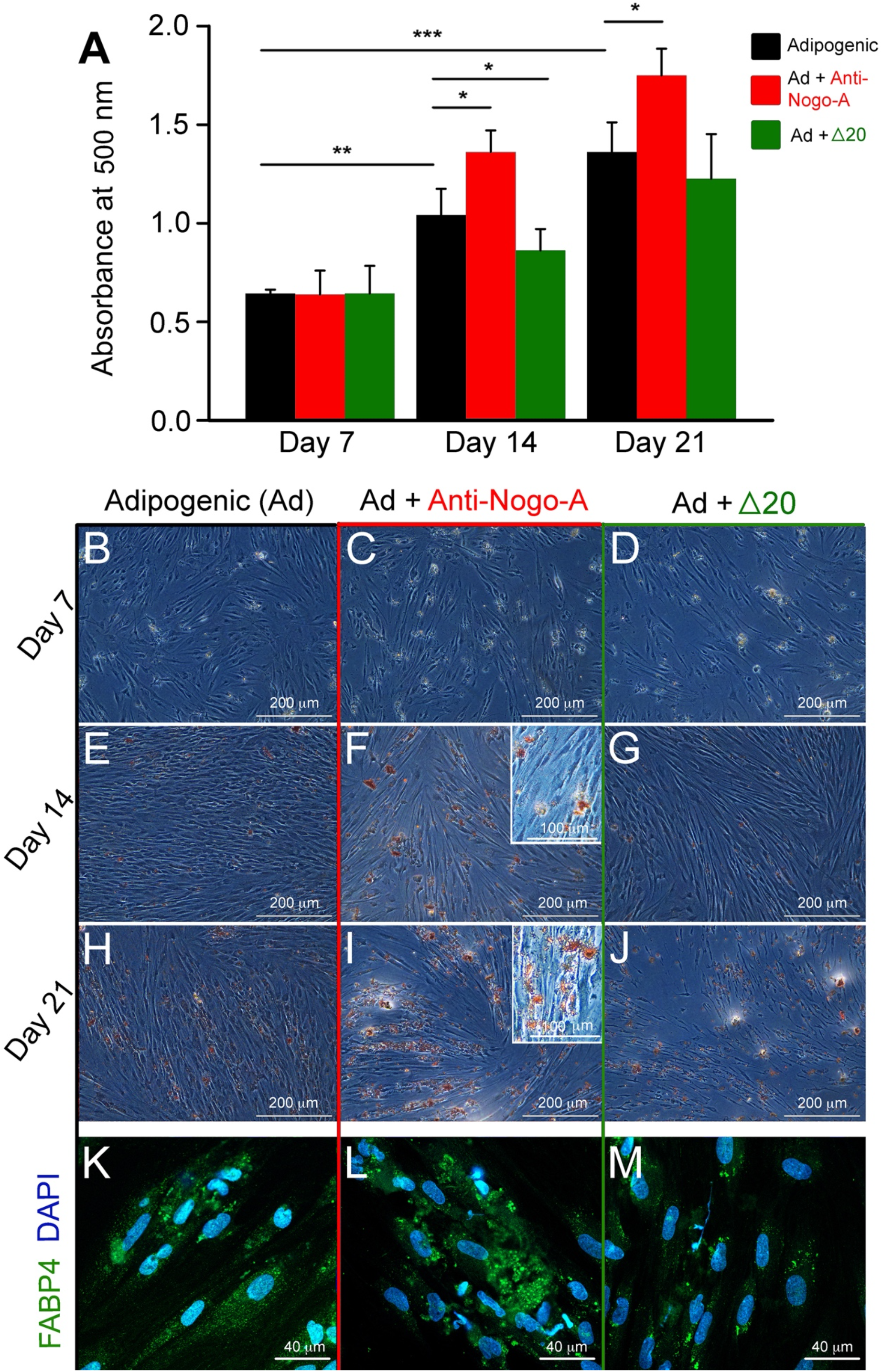
Effects of the Nogo-A protein (Nogo-A Δ20 fragment) and anti-Nogo-A antibody on the adipogenic differentiation of hDPSCs. (A) Quantification of Oil Red O staining in hDPSCs cultured in adipogenic medium (Ad) alone (black colored columns), adipogenic medium supplemented with the anti-Nogo-A antibody (red colored columns), and adipogenic medium supplemented with the Nogo-A Δ20 protein (green colored columns) for 7, 14 and 21 days. Four independent biological replicates were used. The bars represent the means ± standard errors. Asterisks above the error bars denote significant differences (*, p < 0.05; **, p<0.01; ***, p<0.001) by one-way analysis of variance (ANOVA). (B-J) Oil Red O staining indicating the formation of lipid droplets (red color) in hDPSCs cultured in adipogenic medium alone (B, E, H), in adipogenic medium supplemented with the anti-Nogo-A antibody (C, F, I), and in adipogenic medium supplemented with the Nogo-A Δ20 protein (D, G, J) for 7, 14 and 21 days. Inserts in (F) and (I) are higher magnifications demonstrating the formation of lipid droplets (red color) in anti-Nogo-A antibody-treated hDPSCs. (K-M) Fatty Acid Binding Protein 4 (FABP4) immunofluorescence staining (green color) in hDPSCs cultured for 21 days in adipogenic medium alone (K), in adipogenic medium supplemented with the anti-Nogo-A antibody (L), and in adipogenic medium supplemented with the Nogo-A Δ20 protein (M). DAPI (blue color) marks the nuclei of the cells. Scale bars are as indicated.

Higher adipogenesis in hDPSCs from group B was also associated with intense immunofluorescence staining for the fatty acid binding protein 4 (FABP4; Fig. 6L), which was almost absent in hDPSCs from group C as compared to hDPSCs of group A (compare Fig. 6M to Fig. 6K).

RT-qPCR analysis of the *LPL* (Fig. 7C) and *PPARG* (Fig. 7D) adipogenesis-related genes showed significant upregulation of their expression in hDPSCs from group B already at day 7, indicating earlier onset of adipogenesis. High levels of *LPL* and *PPARG* expression in hDPSCs of group B were maintained throughout the 21 days culture period (Fig. 7C, D), corroborating the increased Oil Red O staining in these cultures (Fig. 6F, I). In contrast, *LPL* and *PPARG* mRNA expression in hDPSCs from group C was maintained at low levels throughout the culture period, which indicated delayed onset and inhibition of adipogenesis (Fig. 7C-D).

**Figure 7.**
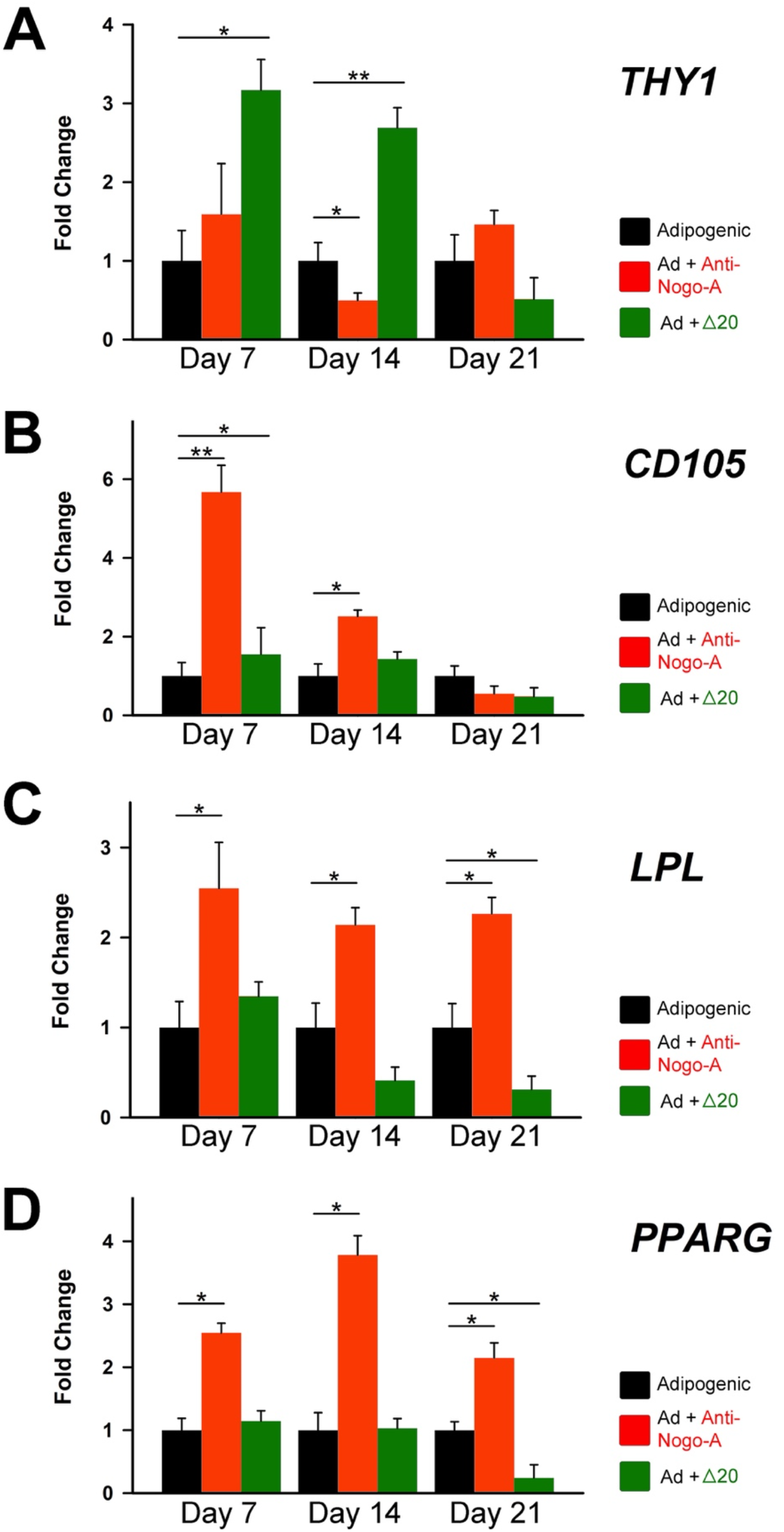
Effects of the Nogo-A protein and anti-Nogo-A antibody on the expression of genes during hDPSC adipogenic differentiation. qRT-PCR analysis of the relative expression levels of *THY1/CD90* (A), *END/CD105* (B), *LPL* (C) and *PPARG* (D) in hDPSCs cultured in adipogenic medium (Ad) alone (black columns), in adipogenic medium supplemented with the anti-Nogo-A-antibody (red colored columns), and in adipogenic medium supplemented with the Nogo-A Δ20 protein (green colored columns) for 7, 14 and 21 days. Values, normalized to GAPDH, shown are the mean ± SD of triplicates. Statistical analysis used Student’s t-test (significant difference at *, p<0.05; **, p<0.01).

#### The anti-Nogo-A antibody accelerates the neurogenic differentiation of hDPSCs

hDPSCs undergo neurogenic differentiation if cultured under appropriate conditions (e.g. Neurobasal A medium supplemented with EGF and FGF) ^13^. Addition of neurogenic medium (control, group A) induces generation of neuron-like cells within 14 days (Fig. 8A), positive for Neurofilement light chain (NFL) and Tubulin Beta-III (TUBB3) staining (Fig. 8D). Neurogenic medium supplemented with the anti-Nogo-A antibody (group B) affected both the cell bodies and the forming axons in differentiating hDPSCs (Fig. 8B), which were also NFL- and TUBB3-positive (Fig. 8E, H). Cell bodies in group B increased in size and presented a rounder, more neuron-like appearance (Fig. 8B), when compared to hDPSCs of group A (compare Fig. 8B with Fig. 8A). A similar increase was observed in the length of their axons and axon-like structures (Fig. 8B). In contrast, hDPSCs cultured in neurogenic medium supplemented with the Nogo-A Δ20 protein (group C) showed modest neurogenic differentiation (Fig. 8C) and less abundant NFL (Fig. 8F) and TUBB3 (Fig. 8I) staining as compared to hDPSCs from group A (compare Fig. 8C, F, I with Fig. 8A, D, G).

**Figure 8.**
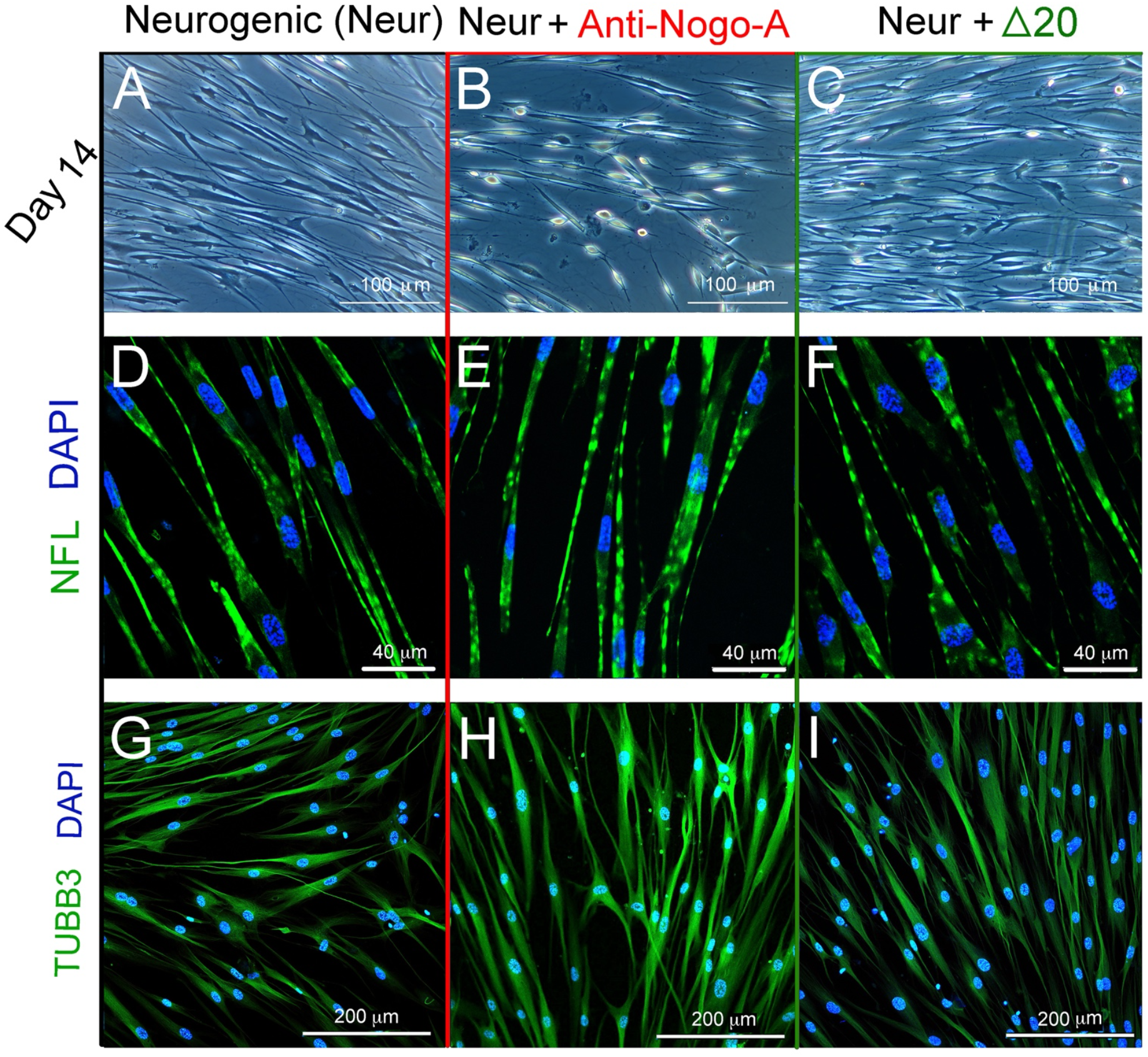
Effects of the Nogo-A protein and anti-Nogo-A antibody on hDPSCs neuronal differentiation. (A-C) Morphological changes of hDPSCs cultured in neurogenic induction medium (Neur) alone (A), in neurogenic medium supplemented with the anti-Nogo-A-antibody (B), and in neurogenic medium supplemented with the Nogo-A Δ20 protein (C) for 14 days. (D-F) Neurofilament-L (NFL) immunofluorescent staining (green color) in hDPSCs cultured for 7 days in neurogenic medium alone (D), in neurogenic medium supplemented with the anti-Nogo-A-antibody (E), and in neurogenic medium supplemented with the Nogo-A Δ20 protein (F). DAPI (blue color) marks cell nuclei. (G-I) Tubulin Beta-III (TUBB3) immunofluorescent staining (green color) in hDPSCs cultured for 7 days in neurogenic medium alone (G), in neurogenic medium supplemented with the anti-Nogo-A-antibody (H), and in neurogenic medium supplemented with the Nogo-A Δ20 protein (I). DAPI in blue color.

These changes were confirmed by RT-qPCR analysis for *Doublecortin* (*DCX*) *Tubulin Beta-III* (*TUBB3*), and *NFL*, which are markers for neuronal progenitors, early neuronal differentiation, and differentiated neurons, respectively. Expression of these genes, as well as that of *CD105*, but not of *THY1*, was significantly upregulated in hDPSCs from group B (Fig. 9A-E). A decrease in the values of *TUBB3* (Fig. 9C) and *NFL* (Fig. 9E) was observed upon 21 days of culture, plausibly because the neuronal identity of the hDPSCs was acquired at earlier stages (i.e. day 14). Interestingly, *THY1* expression was upregulated in hDPSCs of group C, suggesting *THY1*-expressing cells as a potential target of Nogo-A.

**Figure 9.**
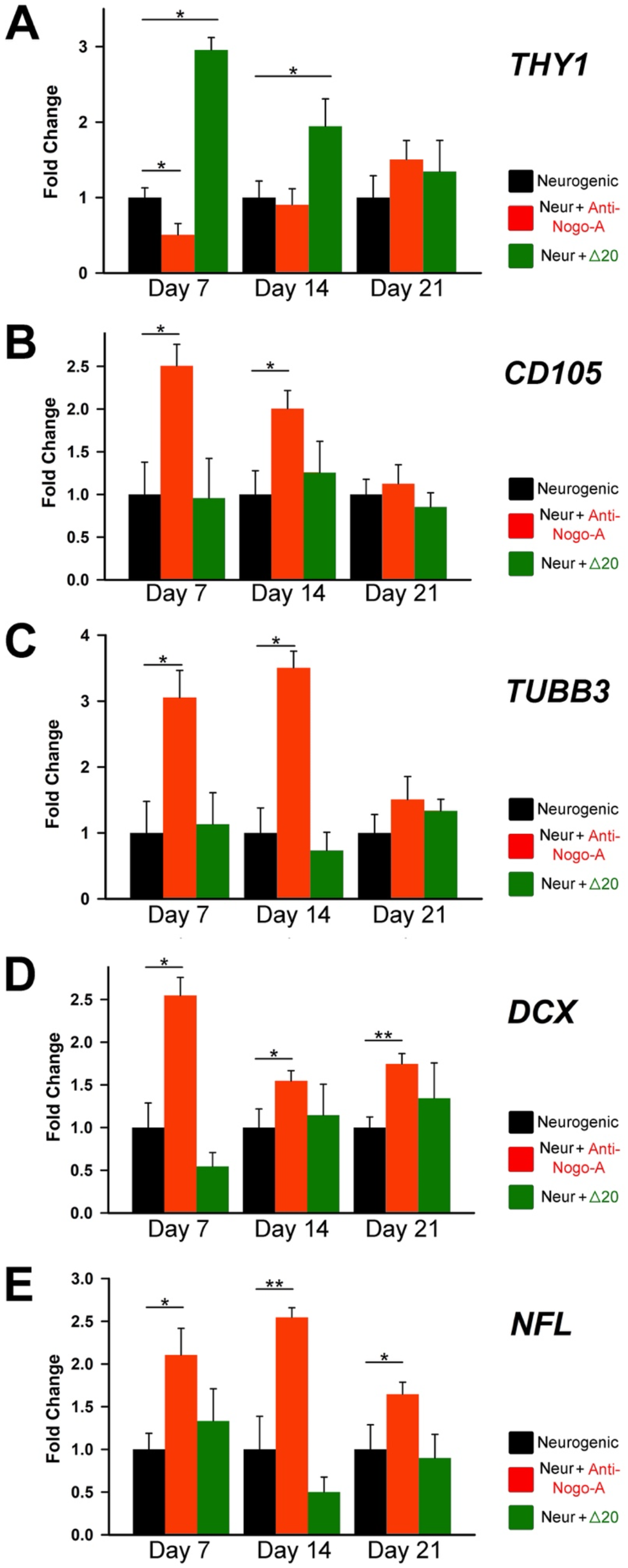
Effects the Nogo-A protein and anti-Nogo-A antibody on the expression of genes during hDPSC neuronal differentiation. qRT-PCR analysis of the relative expression levels of *THY1/CD90* (A), *END/CD105* (B), *TUBB3 (Tubulin Beta-III;* C), *DCX (Doublecortin;* D) and *NFL (Neurofilament-L;* E) in hDPSCs cultured in neurogenic medium (Neur) alone (black columns), in neurogenic medium supplemented with the anti-Nogo-A antibody (red color columns), and in neurogenic medium supplemented with the Nogo-A Δ20 protein (green color columns) for 7, 14 and 21 days. Values, normalized to Glyceraldaeyde-3-Phosphate Dehydrogenase (GAPDH), shown are the mean ± SD of triplicates. Statistical analysis used Student’s t-test (significant difference at *, p<0.05; **, p<0.01).

## Discussion

Regenerative medicine offers alternative therapeutic approaches to the currently employed clinical practices for the repair or replacement of damaged tissues ^16,23,46,78–80^. Translation of experimental regenerative therapies to the patients is one of the biggest challenges of our times, which aim to advance a new medical discipline expecting to tackle major unmet healthcare needs ^80–83^. Regenerative therapies may include both stem cell-based and molecule-based approaches to promote the repair of tissues and restore their structure and functions ^23,82,84^. Mesenchymal stem cells (MSCs) have gained great attention in the past few decades due to their self-renewal, multilineage differentiation potentials, and immunomodulatory properties, which are important features for the efficacy of cell-based regenerative treatments ^82^. Another advantage of MSCs is their fast availability in clinically relevant quantities ^79,81^. Innovative approaches based on MSCs and specific molecules that are entering the clinical arena could also be used in dentistry in order to restore the structural integrity and physiology of teeth ^23,30,38,47,79^. Pioneer *in vitro* and *in vivo* studies have already demonstrated the potential of human dental pulp stem cells (hDPSCs) for the regeneration of both dental and non-dental tissues in patients and in experimental animals ^38,45,85^. However, a key challenge for tissue regenerative treatments is the efficient use of specific molecules, since our knowledge about their way of action in the various cell types and contexts is mainly agnostic ^46,79^. Indeed, most of the information is often based on *in vitro* experimental setups designed for distinct environments that do not necessarily reflect the *in vivo* patient-specific situation ^82,83,86^. Several signaling pathways, including Wnt, Notch, TGF-β, EGF, VEFG and Neurotrophins have been used for controlling stem cell behavior in pathological conditions ^25,87–90^. The mode of action of these molecules can provide insights into their effects and adaptive behavior in specific cells and tissues during regenerative treatments.

Due to the beneficial effects of Nogo-A inactivation for the regeneration of injured nerve fibers ^91–94^, this molecule is the target in clinical trials aiming at the treatment of severe spinal cord lesions ^72^. Antibodies or decoy receptors (‘Nogo-traps’) that block the function of Nogo-A could be tools of high potential for these therapies. Besides its role in neuronal regeneration, Nogo-A is a potential target for enhancing vascular regeneration and functionality, since its inhibition restores vascularization in the central nervous system in Parkinson’s disease ^95^ and upon stroke in mice ^66^. These Nogo-A properties could also be applicable in dentistry for recovering tooth physiology upon injury by regenerating the damaged neuronal and vascular networks within the dental pulp.

Stem cell populations reside in specific niches that maintain stem cells in an undifferentiated status and guide their fates ^3,5,78^. The dental pulp contains several stem cell niches in precise anatomic locations that are composed by fibroblasts, immune cells, vascular, and neuronal structures ^8,11,18–20,89^. Our results show that these dental pulp compartments are characterized by high levels of Nogo-A expression and that the decision whether stem cells will remain in their quiescent, immature state or undergo differentiation is influenced by Nogo-A. Stem cell fate changes and cytodifferentiation are characterized by a shift in gene expression that involves the coordinated action of transcription factors, DNA methylation and chromatin remodeling activities ^96–98^. The cell surface protein Thymocyte Antigen 1 (THY1, also known as CD90) is a marker for MSCs and hDPSCs and plays critical roles in the maintenance of their pluripotency and fate choices ^31,99,100^. Our *in vitro* differentiation assays showed that addition of Nogo-A increases the levels of *THY1* in hDPSCs in all experimental setups tested, suggesting that Nogo-A is critical for maintaining the stemness of hDPSCs and inhibiting their differentiation towards osteocytes, adipocytes, and neurons. In contrast, the anti-Nogo-A antibody significantly enhances hDPSCs differentiation, which is in accordance with previous studies showing increased differentiation potential of MSCs upon reduction of *THY1* expression ^101^. CD105 is the main glycoprotein of human vascular endothelium, which is commonly used as another MSC marker ^10,40,102,103^. Antibody mediated blockade of Nogo-A decreased the levels *CD105* in hDPSCs in all experimental groups. It appears that administration of the anti-Nogo-A antibody has opposite effects in the *THY1*- and *CD105*-expressing hDPSCs, thus differently affecting the fate and differentiation potential of these two stem cell populations ^11^.

Our results highlight important, yet neglected, effects of Nogo-A in mineralized tissue formation that could improve the actual clinical treatments by accelerating the repair of dental or other hard tissues, such as dentin and bone. Enhancing bone regeneration is a big challenge for orthopedic research ^42,81,84^. A faster mineralization allows quicker bone healing that is one of the critical steps in bone, craniofacial and dental regenerative medicine ^16,42,79,84^. Nogo-A exerts an inhibitory role in hDPSCs osteogenic differentiation by reducing the expression of osteogenesis related genes, such as *RUNX2*, *OSX* and *ALP*, thus affecting the timing and the extent of mineralization. Indeed, onset of mineralization is accompanied by increased expression of these genes: *RUNX2* is a major control gene for MSCs differentiation into preosteoblasts ^104–106^, *OSX* is involved in the transition of preosteoblasts to mature osteoblasts ^107,108^, and *ALP* is an early osteogenic gene essential for the mineralization process ^109,110^. Their upregulation in hDPSCs cultured in presence of the anti-Nogo-A antibody correlates with the accelerated and extensive formation of mineralized nodules. While *in vitro* differentiation assays do not faithfully mimic the physiological environment and cannot be yet translated to the clinics, they point to exciting new possibilities for improving hard-tissues repair. Future follow-up *in vivo* studies using small or larger animal models will be very useful in order to obtain more robust conclusions concerning the biological functions of Nogo-A in osteogenesis and bone-tissue regeneration.

The present findings demonstrate that Nogo-A also regulates hDPSCs fate towards adipogenic differentiation. While hDPSCs do not exhibit extensive adipogenesis as other MSC populations (e.g. bone marrow stem cells) ^13,29^, previous studies showed they are able to generate limited number of adipocytes evidenced by increased expression of the adipogenesis genes, including *Peroxisome Proliferator-Activated Receptor-gamma2 (PPAR-γ2*) and *Lipoprotein Lipase* (*LPL*), as well as increased Oil Red O staining detecting intracellular lipid droplets ^29^. The transcription factor PPAR-γ2 is a master regulator of adipogenesis and glycose/lipid metabolism ^111,112^ and LPL plays a central role in lipid metabolism, lipid transport, and triglycerides concentration in preadipocytes ^113,114^. Addition of the anti-Nogo-A antibody significantly upregulated *LPL* and *PPAR-γ2* expression and enhanced Oil Red O staining in hDPSCs cultured in adipogenic conditions, although the shift shape of hDPSCs from fibroblastic to spherical that characterizes adipogenic differentiation is not observed ^115^.

The neurogenic potential of hDPSCs is assessed by the increased expression of the neuronal genes *Doublecortin (DCX), Neurofilament light chain (NFL*) and *Tubulin β-III*. DCX is important for modulating and stabilizing microtubules to ensure effective migration of neuronal progenitor cells ^116,117^, NFL is a structural protein forming neurofilaments and plays a role in the regulation of synaptic transmission and organelle trafficking ^118^, and *Tubulin β-III* expression could be a common feature of stem cells and neuronal cells ^119^, and confirms hMSCs differentiation towards neurons ^120^. Addition of the anti-Nogo-A antibody upregulated *DCX, NFL* and *Tubulin β-III* expression in hDPSCs cultured in neurogenic medium. Previous studies have shown that upregulation of these genes correlates with accelerated neuronal outgrowth ^51,52,91,121^. A small interfering RNA (siRNA) technique has been explored both *in vivo* and *in vitro* to specifically suppress Nogo-A to promote axonal growth and repair in a multiple sclerosis murine model ^122^. However, the efficiency of the Nogo-A silencing was only 75% in cultures of an established cell line (N2A), suggesting that the efficiency of siRNA Nogo-A in our experimental setup is likely to be even lower, since we used hDPSCs from primary cultures. Although a confirmation of our results by a siRNA knock down experiment would be interesting, a negative result would not be telling due to potentially insufficient knock down.

In conclusion, our results show that Nogo-A is expressed in the various cell populations composing the dental pulp of human teeth, including odontoblasts, fibroblasts, cells of the vascular and neuronal networks and hDPSCs. The present *in vitro* findings demonstrate that Nogo-A plays an important role in the fate choices and multilineage differentiation potential of hDPSCs by maintaining their stemness and regulating their differentiation towards adipocytes, osteocytes, and neurons (Fig. 10). Inhibition of its function results in faster cell differentiation events, pointing to Nogo-A as a suitable candidate molecule for future regenerative therapies in the clinics, in order to restore tissue integrity and function.

**Figure 10.**
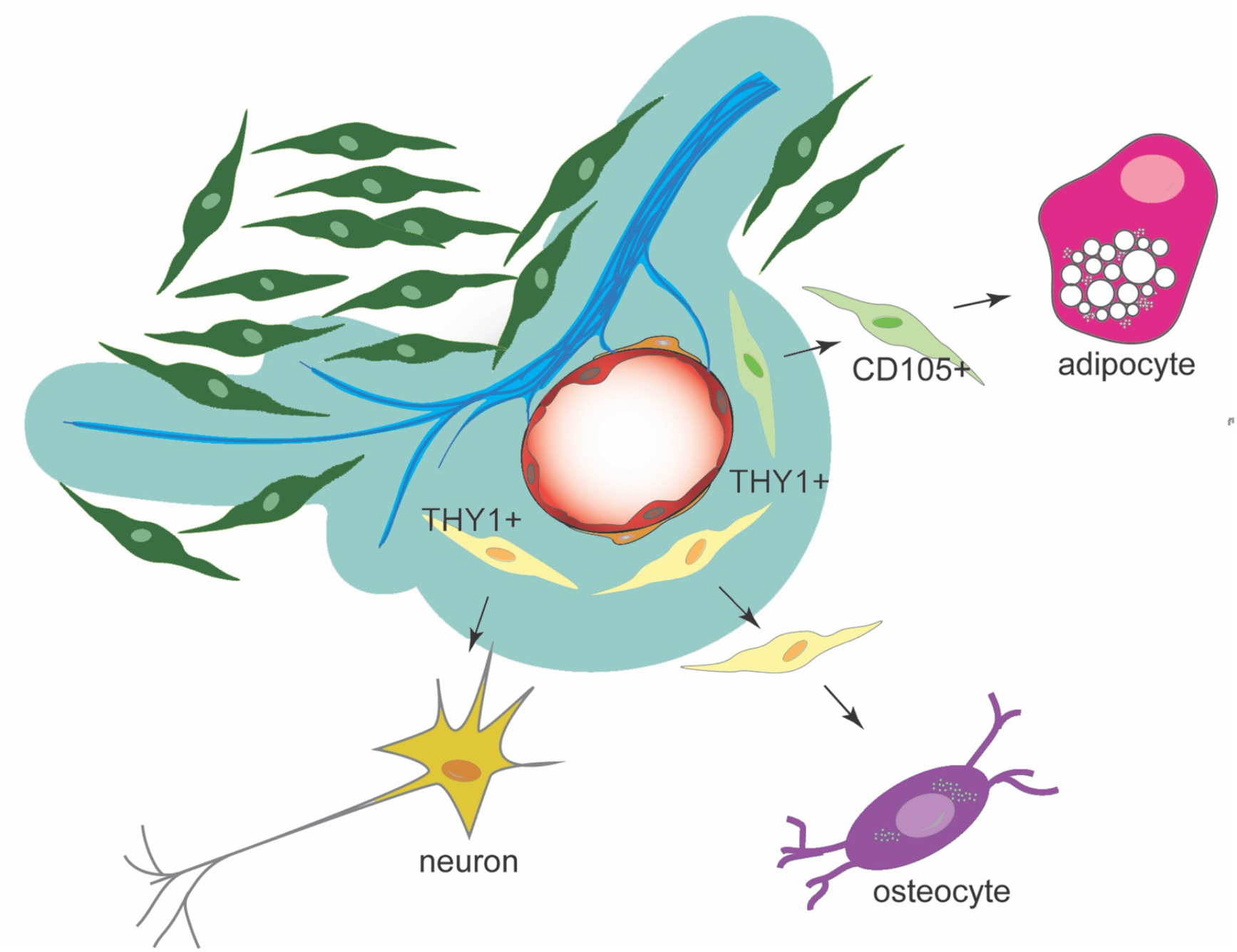
Cartoon depicting the effects of Nogo-A and anti-Nogo-A-antibody on the hDPSC differentiation into adipogenic, osteogenic and neurogenic lineages. Based on our hypothesis, in osteogenic and neurogenic microenvironments, hDPSCs derived from the THY1-positive cell lineage (yellow color) enhance their differentiation potential towards either osteoblasts or neurons in absence of Nogo-A. In adipogenic microenvironment, hDPSCs originated from the CD105-positive cells (light green color) are directed towards an adipocytic fate in the absence of Nogo-A. Dental pulp fibroblasts in dark green color, nerves in blue color, vessels in red color. The territory depicted with the blue/green color indicates areas with various levels of Nogo-A protein expression.

## Acknowledgments

We thank Dr Michael Maurer and Dr Zorica Ristic (Institute for Regenerative Medicine, University of Zurich, Schlieren) for providing the anti-Nogo-A blocking antibody (11C7) and the recombinant Nogo-A- 20 protein fragment.

## Author Contributions

Conceptualization: T.A.M. Methodology: T.A.M, C.F.L., J.C. Data analysis: T.A.M., C.F.L., J.C., A.B., P.P. Validation: T.A.M., C.F.L., J.C., M.E.S., A.B., P.P. Formal analysis: T.A.M., C.F.L., J.C., A.B. Investigation: C.F.L., J.C. Resources: T.A.M., M.E.S. Data curation: T.A.M., C.F.L., J.C., M.E.S., P.P. Writing – original draft: T.A.M., C.F.L., A.B. Writing – Review & Editing: T.A.M., C.F.L., J.C., M.E.S., A.B. Visualization: T.A.M., C.F.L., J.C., M.E.S., A.B., P.P. Supervision: T.A.M. Project administration: T.A.M. Funding acquisition: T.A.M., M.E.S., P.P.

## Funding

This work was financially supported by institutional funds from the University of Zurich (UZH), the Swiss National Science Foundation (SNSF; grant number 31003A_179389), and the Swiss Dental Association (SSO; grant number 313-19).

## Institutional Review Board Statement

No ethical approval required for this study.

## Informed Consent Statement

Informed consent was obtained from all subjects involved in the study, as mentioned in the materials and methods section

## Data Availability Statement

Not applicable.

## Conflicts of Interest

The authors declare no conflict of interest.

## Notes

### Competing Interest Statement

The authors have declared no competing interest.

